# comparison of genetic diversity and phylogenetic structure of 18S gene of *Trichomonas vaginalis* in Hainan and Xinjiang provinces

**DOI:** 10.1101/2024.07.04.602107

**Authors:** Jun Liu, Xiaoyang Dong, Daping Wang, Min Chen, Jing Nie, Man Xiao

## Abstract

*Trichomonas vaginalis* is a kind of flagellate parasite endemic all of the world. study compared the difference of genetic diversity and phylogenetic structure of 18S ribosomal RNA gene of *T. vaginalis* from Hainan and Xinjiang Provinces, China. 20 samples and 47 samples from Hainan and Xinjiang respectively which confirmed for infection with T.vaginalis. The sequences were aligned using MEGA 11 software and phylogenetic trees were drawn by Neighbor-Joining method. The number of mutation, nucleotide diversity, and haploid diversity were analyzed using Dnaspv5 software. For Hainan samples, the analyses showed 31 polymorphisms, creat different haplotypes with a haplotype diversity of 0.589. Nucleotide diversity and average nucleotide different among *T*.*vaginalis* were estimated as 0.00679 and 3.447, respectively.Tajima’s D value in Tajiama’s neutrality test was -2.48147,which was significant (p<0.001). But for Xinjiang samples, the analyses showed 31 polymorphisms, creat different haplotypes with a haplotype diversity of 0.731. Nucleotide diversity and average nucleotide different among T.vaginalis were estimated as 0.00497 and 2.50435, respectively. Tajima’s D value in Tajiama’s neutrality test was -2.26284, which was significant (p<0.01). For all the sequence in this study, all the sequences form one phylogenetic tree, some of the sequence from two provinces showed a certain aggregating phenomenon. The differences between the samples from two province maybe for the local environment, ethnic, sample number and so on, it need further study for T.vaginalis.

## 1. Introduction

*Trichomonas vaginalis* (T. vaginalis) is the pathogen of trichomoniasis, which is the most common sexually transmitted non-viral infection around the world [1], its surpasses the total of syphilis, chlamydia and gonorrhea, it has over 276 million cases annually[2,3]. T.vaginalis is a kind of flagellated protozoan, which resides exclusively in the vaginal, urethra, epodidymis and prostate gland of human. Despite the case of trichomoniasis was large, T.vaginalis still a highly neglected pathogen and poorly studies[4].Clinical manifestations of trichomoniasis range from human urogenital tract inflammation to infertility, adverse pregnancy outcomes such as low infant birth weight and preterm delivery[5,6].

Except using wet mount microscopy, staining techniques, several methods also have used for detecting trichomoniasis, including culture, immunodiagnosis, and molecular diagnosis.[7,8,9,10].Kinds of genes or tools have used to study teh genetic properties of T.vaginalis, including actin gene, RFLP, ITS, qRT-PCR, PCR sequencing and so on[11,12,13,14]. Several researchers using microsatellites and single-copy genes acted polymorphic makers to study genetic diversity in T.vaginalis, and has been mainly identified two population[15]. Some studies have been done on the relationship between the clinical symptoms of trichomoniasis and its genotype, and the genotype of T.vaginalis with the drug susceptibility and symbiosis[16].

## 2. Materials and Methods

### 2.1. Sample collection

There were sixty-seven sample from women (range 18-61 years) who infected with T.vaginalis in total, twenty (range 18-61years) from the First Affiliated Hospital of Hainan Medical University, and forty-seven (range 20-52 years) from the First Affiliated Hospital of Xinjiang Medical University.

### 2.2. Sample evaluation by microscopy and genomic DNA extraction

Combined with clinical manifestations and testing techniques, all samples confirmed infected with T.vaginalis. Then the samples were sent quickly to the laboratory. Strictly followed the protocol, extracted total genomic DNA from all samples. The DNA stored at -20°C for future testing.

### 2.3. PCR amplification and sequencing

To amplify the 18S ribosomal genes in T.vaginalis, using the same specific primers, amplification system, and procedure as used before[17]. After PCR amplification, the products were electrophoresis on a 2% agarose gel in TAE buffer at 100 volts. The gel was stained with GeneRed and visualized under ultraviolet transilluminator to make sure the size of amplified product was 582 bp by comparison with a commercial 100 bp DNA ladder (Tiangen).

### 2.4. Gene analysis

After gene sequencing, by BLAST on NCBI to verity the sequences from T.vaginalis. Then using Clustal W algorithm in BioEdit 7.0.5.3[18] to aligning multiple sequences. The sequences were aligned using MEGA 11 software[19] and phylogenetic trees were drawn by Neighbor-Joining method. Using DnaSP5 software to computed the haplotype diversity and nucleotide diversity and insight the genetic variability of these gene sequences.

## 3. Results

After isolation of DNA from fresh specimens, segments of the 18S ribosomal gene sequences belonging to *Trichomonas vaginalis* were successfully amplified and underwent sequencing procedures. All samples from 20 Hainan females, 20 T. vaginalis 18S ribosomal RNA sequences were identified (labeled HN-1 through HN-20) and 47 from Xinjiang female (labeled XJ-1 through XJ-47). To construct a rooted phylogenetic network, the 67 sequences identified in this study and another 12 reference sequences reported in GeneBank were included, which were from American(U17510), Philippines (JX943583 and KM282377), China Henan province (KM603320, KM603325, KM603331, KM603336, KM603339, KM603343, KM603347), Iran (KX061402, KX272849).

### 3.1 Genetic variation

All samples were amplified 18S rRNA gene successfully, the length of sequence ranging from 544 to 623 bp. After trimming, the final sequence alignment contained 79 sequences, 515 bp positions. They contain 51.3% G+C bases. A total of 31 polymorphisms for samples from two provinces, but for samples from Hainan, 8 haplotypes were identified, while 10 haplotypes for Xinjiang samples. Each population displayed very low nucleotide diversity. The average number of nucleotide differences for two location samples were inconformity. Both of two sources samples, there were significant negative value of Tajima’s D (table 1).

**Table 1.** Diversity of 18S ribosomal RNA gene sequences in *T. vaginalis* from Hainan and Xinjiang.

|  | Hainan | Xinjiang |
| --- | --- | --- |
| Number of sequences | 20 | 47 |
| Number of Haplotypes | 8 | 10 |
| Haplotype (gene) diversity | 0.589±0.130 | 0.731±0.044 |
| Number of polymorphic | 31 | 31 |
| Nucleotide diversity, Pi | 0.00679±0.00458 | 0.00497±0.0016 |
| Average number of nucleotide differences | 3.447 | 2.50435 |
| G+C content | 0.513 | 0.513 |
| Tajima's D | -2.48147 | -2.26284 |
| Statistical significance | $P < 0.001$ | $P < 0.01$ |

### 3.2 Phylogenetic diversity

All sequences form a big evolutionary tree, analyzed from the whole, the gene sequences from Hainan are more primitive in evolution than from Xinjiang. And the sequences from Xinjiang have a wide range of evolutionary manifestations, some of them form a evolved higher-order communities. all the reference sequences are located between two location samples in the evolutionary tree (fig 1).

**Fig 1.**
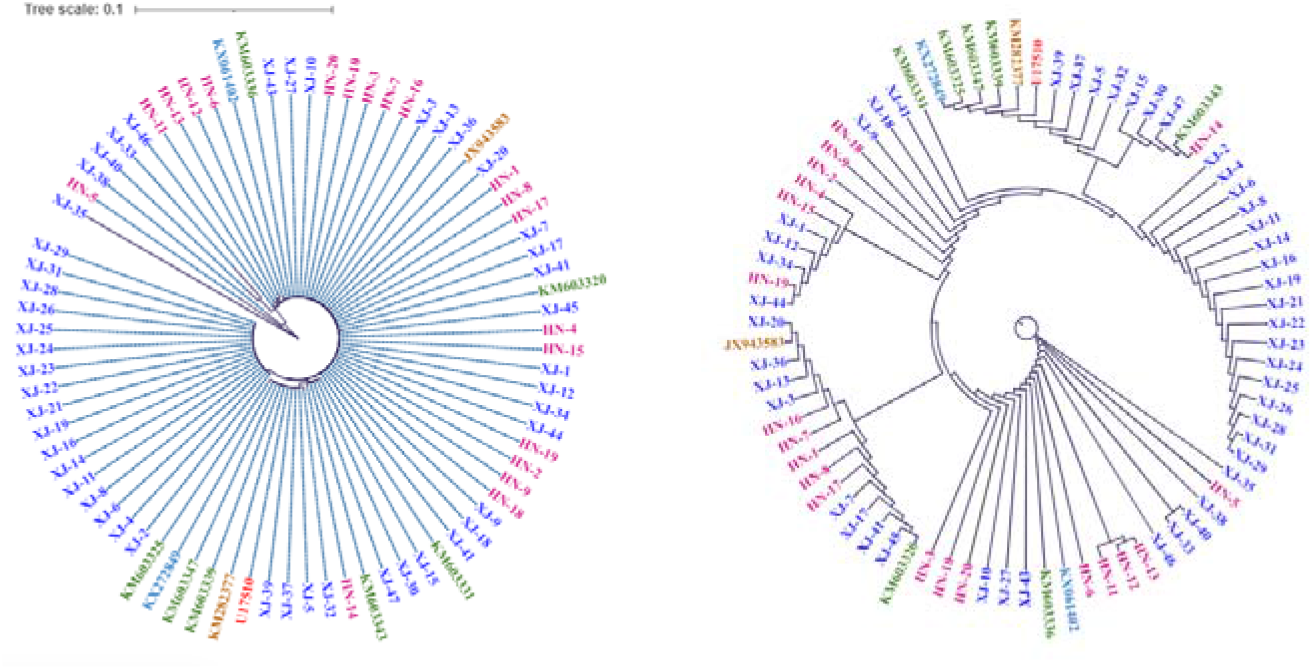
The evolutionary tree of partial 18S ribosomal RNA gene sequences of T.vaginalis Using different colour to show the samples from different regions.

## 3. Discussion

Although the number of people infected by *T*.*vaginalis* is large, the importance of its research is not proportional to the number of infected people.It is the first time to report the genetic diversity in *T*.*vaginalis* in Hainan, China. Even though there have been some international studies in USA, Philippine, Iran, Australia and Ghana [11,13,15,20], and Henan, Xinjiang, Gansu province in China[17,21,]. In this study, Hainan *T*.*vaginalis* isolates showed a greater degree of negative selection, suggest that Hainan’s unique geographical location, there has been little movement between humans since ancient times, resulting in local organisms showing a certain regional characteristics.

In this study, the average nucleotide diversity in Hainan higher than that found among T. vaginalis isolates in a report (0.0035 difference/site) [15],but lower than that of Xinjiang. DNA polymorphism analysis revealed high genetic diversity of T. vaginalis isolates. Although moderate-to-high genetic differentiation between isolates from different sites was observed, T. vaginalis isolates from different provinces should be taken as a whole to research. Our data support demographic expansion of the T. vaginalis population.

## Acknowledgements

This work was supported by the Hainan Provincial Natural Science Foundation of China (820QN274) and Hainan University Scientific Research Project (Hnky2020-38).

## Contributors

All named authors contributed to this article.

## Conflict of interest

The authors declare that they have no competing interests.

## Ethics approval

Human Ethical Committees of Hainan Medical University, and permission from Department of Obstetrics and Gynecology, the First Affiliated Hospital of Hainan Medical University and the the First Affiliated Hospital of Xinjiang Medical University, the Department of Gynecology, Hainan Children’s Hospital.

## Notes

### Competing Interest Statement

The authors have declared no competing interest.

## References

[1] Schwebke JR, Burgess D. Trichomoniasis. Clin Microbiol Rev. 2004; 17(4): 794–803.

[2] World Health Organization. Report on global sexually transmitted infection surveillance. 2018.1–74.

[3] Rowley J, Hoorn SV, Korenromp E, et al. Chlamydia, gonorrhoea, trichomoniasis and syphilis: Global prevalence and incidence estimates, 2016. Bull World Health Organ. 2019; 97(8):548–562.

[4] Alexandra IE, Juan Jose Nogal-Ruiz. The past, present, and the future in the diagnosis of a neglected sexually transmitted infection: trichomoniasis. Pathogens. 2024,13(2):126.

[5] Petrin D, Delgaty K, Bhatt R, et al. Clinical and microbiological aspects of Trichomonas vaginalis. Clin Microbiol Rev. 1998; 11(2):300–317.

[6] Cotch MF, Pastorek JG, Nugent RP, et al. Trichomonas vaginalis associated with low birth weight and preterm delivery. Sex Transm Dis. 1997; 24(6):353–360.

[7] Hobbs MM, Sena AC. Modern diagnosis of Trichomonas vaginalis infection. Sex. Transm. Infect. 2013, 89, 434–438.

[8] Khatoon R, Jahan N, Khan HM, et al. Evaluation of different staining techniques in the diagnosis ofTrichomonas vaginalis infection in females of reproductive age group. J. Clin Diagn Res. 2014, 8,5–8.

[9] Alderete JF. Advancing prevention of STIs by developing specific serodiagnostic targets: Trichomonas vaginalis as a model. Int. J.Environ. Res. Public Health 2020, 17, 5783.

[10] Bui, HTV, Bui, HT, Chu, SV, et al. Simultaneous real-time PCR detection of nine prevalent sexually transmitted infections using a predesigned double-quenched TaqMan probe panel. PLoSONE 2023, 18, e0282439

[11] Alikhani M, Saberi R, Hosseini SA, et al. Identificaiton of Trichomonas vaginalis genotypes using by actin gene and molecular based methods in southwest of Iran. Rep Biochem Mol Biol. 2021,10(1):135–143.

[12] Demirag S, Malatyali E, Ertug S, et al. Determination of trichomonas vaginalis genotypes using PCR-restriction fragment length polymorphism (RFLP). Turkiye Parazitol Derg. 2017;41(4):188–191.

[13] Ertabaklar H, Ertug S, Caliskan So, et al. Use of internal transcribed spacer sequence polymorphisms as a method for trichomonas vaginalis genotyping. Turkiye Parazitol Derg. 2018;42(1):6–10.

[14] Ahmadi MH, Amirizadeh N, Rabiee M, et al. Noninvasive fetal sex determination by realtime PCR and TaqMan probes. Rep Biochem Mol Biol. 2020;9(3):315–323.

[15] Conrad MD, Gorman AW, Schillinger JA, et al. Extensive genetic diversity, unique population structure and evidence of genetic exchange in the sexually transmitted parasite Trichomonas vaginalis. Plos Neglect Trop D. 2012; 6:e1573.

[16] Hampl V, Vanacova S, Kulda J, et al. Concordance between genetic relatedness and phenotypic similarities of Trichomonas vaginalis strains. BMC Evol Biol. 2001;

[17] Jun Liu, Meng Feng, Xiaolan Wang, et al. Unique Trichomonas vaginalis gene sequences identified in multinational regions of Northwest China. BioScience Trends, 2017,11(3);303–307.

[18] Hall TA. BioEdit: a user-friendly biological sequence alignment editor and analysis program for Windows 95/98/NT. Nucleic Acids Symp Ser. 1999, 44:95–98.

[19] Tamura K, Stecher G, Kumar S. MEGA11: molecular evolutionary genetics analysis version11. Mol Biol Evol. 2021,38:3022–3027.

[20] Daniel SS, Alan JL, Jennifer W, et al. Population structure and genetic diversity of Trichomonas vaginalis clinical isolates in Australia and Ghana. Infection, Genetics and Evolutions. 2020, 82:104318.

[21] Meng Mao, Huili, Liu. Genetic diversity of Trichomonas vaginalis clinical isolates from Henan province in central China. Pathogens and Global Health. 2015, 109(5):242–246.

[22] shuhui Xu, Zhixin Wang, Hang Zhou, et al. High co-infection rate of Trichomonas vaginalis and candidatus Mycoplasma Girerdii in Gansu Province, China. Healthcare. 2021,9:206.

